# Discordant calls across genotype discovery approaches elucidate variants with systematic errors

**DOI:** 10.1101/2022.03.24.485707

**Authors:** Elizabeth G. Atkinson, Mykyta Artomov, Konrad J. Karczewski, Alexander A. Loboda, Heidi L. Rehm, Daniel G. MacArthur, Benjamin M. Neale, Mark J. Daly

## Abstract

Large-scale next-generation sequencing datasets have been transformative for informing clinical variant interpretation and as reference panels for statistical and population genetic efforts. While such resources are often treated as ground truth, we find that in widely used reference datasets such as the Genome Aggregation Database (gnomAD), some variants pass gold standard filters yet are systematically different in their genotype calls across genotype discovery approaches. The inclusion of such discordant sites in study designs involving multiple genotype discovery strategies could bias results and lead to false-positive hits in association studies due to technological artifacts rather than a true relationship to the phenotype. Here, we describe this phenomenon of discordant genotype calls across genotype discovery approaches, characterize the error mode of wrong calls, provide a blacklist of discordant sites identified in gnomAD that should be treated with caution in analyses, and present a metric and machine learning classifier trained on gnomAD data to identify likely discordant variants in other datasets. We find that different genotype discovery approaches have different sets of variants at which this problem occurs but that there are characteristic variant features that can be used to predict discordant behavior. Discordant sites are largely shared across ancestry groups, though different populations are powered for discovery of different variants. We find that the most common error mode is that of a variant being heterozygous for one approach and homozygous for the other, with heterozygous in the genomes and homozygous reference in the exomes making up the majority of miscalls.

## Introduction

While next-generation sequencing (NGS) technologies have been transformative for genomics research, they have an appreciable error rate (Ma et al. 2019) as a cost of their high-throughput capacity. To account for this, sophisticated pipelines have been developed for the detection and removal of incorrect sequencing calls (Li et al. 2019; McKenna et al. 2010; Adelson et al. 2019; Highnam et al. 2015; Anderson et al. 2010; Lam et al. 2019). However, even with gold-standard filtering, spurious genotype calls can infiltrate datasets and potentially skew results. This is of particular importance with datasets that aggregate calls generated by multiple genotype discovery approaches, as different strategies have distinct error modes. Identifying variants that have technical artifacts affecting genotype calls is of major importance, as such loci give misleading information regarding population allele frequencies, and could be incorrectly identified as being phenotypically meaningful in gene discovery.

By leveraging the unprecedented size and depth of the gnomAD database (Karczewski et al. 2020; Lek et al. 2016), we comprehensively characterize trends in genotype calling depending on the sequencing technology. Specifically, we note that a subset of variants, despite passing standard quality filters (Karczewski 2017), produce discordant allele frequencies in data generated using different genotype discovery approaches, stemming from unreliable variant calling. This can not be explained by population stratification, as this effect is observed even when looking at the same set of individuals. Such unreliably genotyped variants should therefore be screened out of analyses. Including these variants in gene discovery efforts, particularly in study designs where case and control data are represented by different combinations of sequencing platforms or genotype discovery approaches, could result in their appearance as false-positive associations.

In this article, we comprehensively characterize the observation of discordant genotyping depending on genotype discovery approach using a large set of diverse individuals from the gnomAD database, including a subset of participants who underwent both whole-exome and whole-genome sequencing (WES/WGS). We then validate our findings in two external datasets for which data from multiple genotype discovery approaches are available: the Thousand Genomes Project and the All of Us Research Project (Auton and Salcedo 2015; The All of Us Research Program Investigators 2019). Correcting for this technical error, whether by removing the gnomAD blacklist variants provided here or by identifying user-identified spurious calls with our freely distributed machine learning predictor, should be incorporated as a step in QC pipelines to avoid spurious associations, particularly in large-scale studies aggregating data from multiple sources.

## Results

### Discordance in genotype calls across genotype discovery approaches replicates across ancestries

We sought to compare allele frequencies across variants found within exome and genome sequencing datasets in gnomAD to test whether there are regions with significant bias associated with genotype discovery approach. We first focused on the largest population represented in gnomAD 0.2 – the Non-Finnish Europeans (NFE). Using the full release of gnomAD version 2.1.1, we filtered the data to include only sites that were present and had a quality determination of PASS in both the genomes and exomes (Karczewski et al. 2020). To ensure sufficient power, we filtered for sites with allele count (AC) >10 and ran a Fisher’s Exact Test (FET) on the difference in the number of alternate AC to total alleles (allele number, AN) between these two datasets. A non-negligible fraction of sites were significantly discordant in their calls (**Figure 1A**). We also tested other less stringent AC thresholds (AC>1 and AC>5) and observed that the trends of discordance between the genotype discovery approaches were consistent across AC cutoffs (**Supplemental Figure S**1).

When comparing allele frequencies between sequencing strategies it is critical to control for ancestry, as populations will have differing frequencies at many loci simply due to demography (Auton and Salcedo 2015; Gravel et al. 2011; Bergström et al. 2020). To assess if ancestry affected discordance rates, we ran FET concordance tests across all gnomAD continental groups (**Figure 1B**). The total discordant variant counts were directly related to the sample size of the population in question and were not enriched for any given ancestry (**Supplemental Figure S2**). While novel discordant variants were discovered in each population, most variants observed in other populations were shared with the NFE (86.9%, **Figure 1B, Supplemental Table S1**). Replication of the same variants across multiple ancestries strengthens the argument of a shared technical artifact and suggests that ancestral bias between exome and genome datasets is unlikely to be a confounding factor. Future work may wish to investigate other biological and non-biological factors for an impact on discordance.

**Figure 1.**
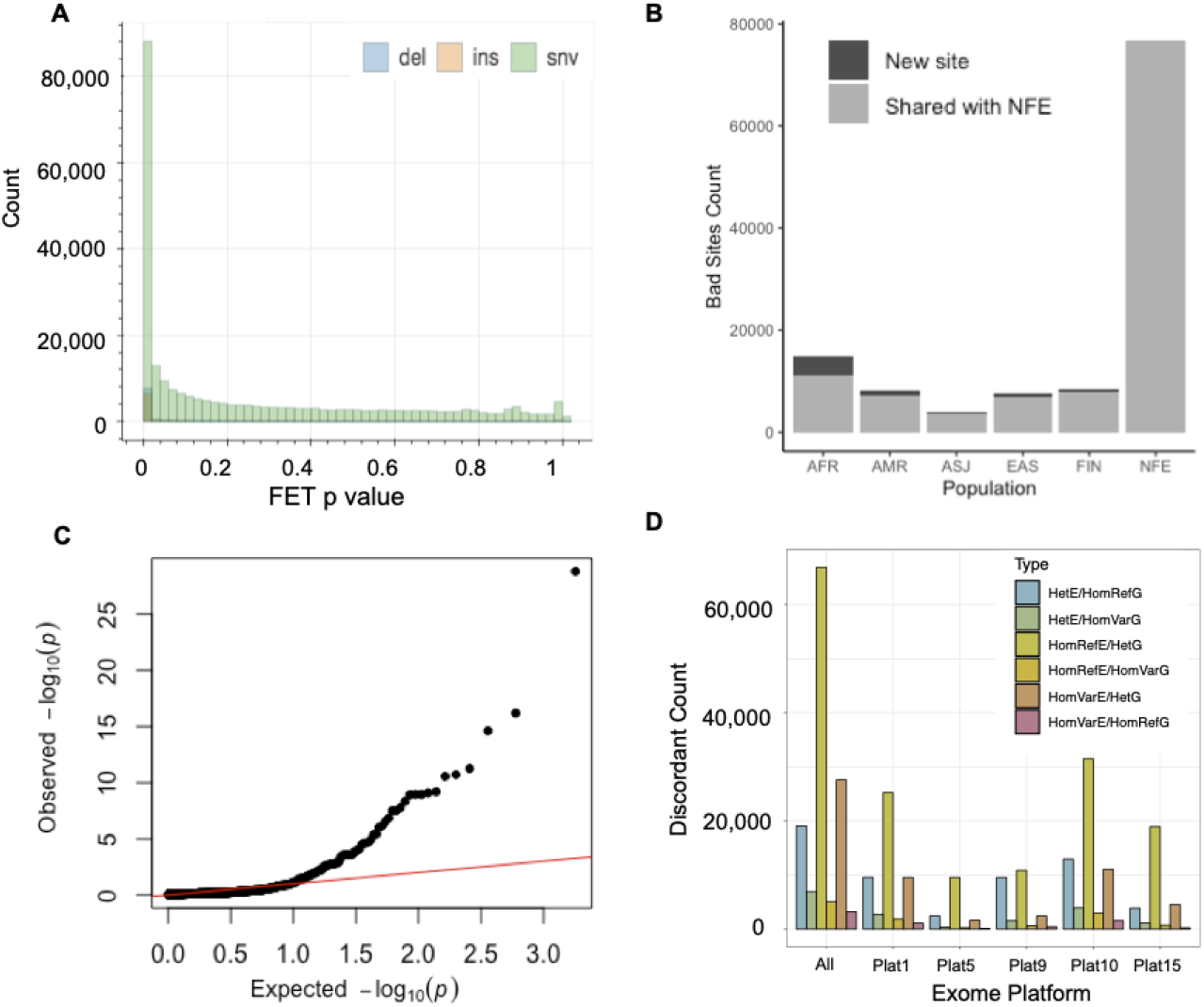
Discordance in genotype calls across NGS genotype discovery approaches. **A**) FET concordance test *p* value for shared, PASS sites in the gnomAD NFE exomes and genomes. Bars are colored by variant type: insertion (ins), deletion (del), or single-nucleotide variant (snv). **B**) ‘Bad’ sites are replicated across ancestry groups in gnomAD. Sites flagged as discordant in both the NFE and another ancestry group are plotted in grey, those new sites not in the NFE are shown in black. **C**) QQ plot for the FET *p* value of shared variants in a set of 946 individuals for whom both WES and WGS data was available. **D**) Different exome captures’ contribution to discordant sites. Bars are colored by the error mode that was observed for the discordant genotype call: heterozygous (het), homozygous reference (HomRef) or homozygous variant (HomVar) in either the exomes (E) or genomes (G).

### Error mode of discordant calls

Next, we aimed to classify the error mode that results in discordant calls using individuals with both WES and WGS data. As 946 gnomAD individuals underwent both whole exome and whole genome sequencing, we were able to examine the rates and error modes of discordant genotype calls without concern over population structure or differing sample composition; since these are the same individuals, any difference in AF and/or genotype calls can be conclusively determined to be due to technical artifacts. We tallied the number of sites in each pairwise exomes-genomes genotype category (6,333 sites; **Supplemental Figure S**3) to classify miscall error mode. Of the 6 possible error modes (homozygous reference/heterozygous, homozygous reference/homozygous variant, or homozygous variant/heterozygous for both dataset directions), we find that the majority of calls, 57.7%, are a heterozygous genotype call in the genomes but a homozygous reference genotype call in the exomes (**Figure 1D, Supplemental Figure S4**). We also note that different sequencing platforms have different rates of discordant calls, though due to sharing restrictions we can not identify platform names with certainty. Overall, ∼16% of the variants that were present and PASS in both exomes and genomes in the overlapping individuals had at least one discordant call.

Next, we examined the discordance of allele frequencies, again with a FET. Usually in cohort-based comparisons, the expected distribution of FET *p value*s is represented by a uniform distribution. Since we are looking at the same individuals, it is expected that allele frequencies should be identical (i.e. *p*=1). We observe the presence of many variants substantially deviating from expectations, representing loci with significantly different MAF in the exomes vs the genomes (**Figure 1C**).

### Identification of discordant sites

We next sought to identify problematic sites failing a FET test of concordant WGS/WES frequency estimates in the largest subset of gnomAD, the NFE. Based on the distribution of *p* values from this test, we decided upon a threshold of *p*<1×10^−5^ to determine the classification of a variant as ‘bad’ or ‘good’ (**Figure 1, Supplemental Figure S5**). Of the 283,287 PASS/PASS variants tested with MAF>0.01 and AC>10, 51,255 (18.1%) failed the FET test and were deemed ‘bad’ while 231,631 (81.8%) passed and were deemed ‘good’. Distributions of metadata features for the good vs bad sites do show trends in several features, though no feature alone perfectly explains the phenomenon (**Supplemental Figure S6**). They also highlight a difference in discordance patterns of indels vs SNVs (**Supplemental Figure S7**). Specifically, SNVs show a pattern of higher AF in genomes as compared to exomes, while indels do not have this trend. Indels are also generally less stable in allele frequency estimates than SNVs. It therefore appears that two distinct technical error modes might be affecting miscalls in indels vs SNVs, rather than one shared mechanism. As many of these indels fell in the low complexity regions of the genome, it is likely that a mapping issue is responsible for their miscalls. A comprehensive description of gnomAD structural variant calling and considerations is published and can be found in a gnomAD blog post (Collins et al. 2020; https://gnomad.broadinstitute.org/news/2019-03-structural-variants-in-gnomad/). To correct this, we therefore recommend excluding the low complexity regions from stringent analyses. In general, when there was discordance, the genomes were found to have a higher MAF than the exomes (**Supplemental Figure S1B**). The trend in MAF difference aligns with the most commonly observed error mode in genotyping.

Having confirmed that there was a systematic and significant AF discordance between genotype discovery approach, we used our FET tests to generate a list of sites harboring this technical artifact that may be excluded from analyses. Again, these blacklisted sites represent variants that were a PASS in gnomAD QC in both the exomes and genomes, but are unreliably genotyped depending on the sequencing technology used. Given a situation where, for example, a case cohort has been exome sequenced while the control cohort has been genome sequenced, such sites could give false-positive associations due to the resulting AF differences. We, therefore, recommend for them to be treated with caution or broadly excluded (in addition to standard cohort QC) unless thorough confirmation of their validity in a particular dataset has been done.

Our analysis of variant allele frequency discordance reflects technical differences between whole exome and whole genome next-generation sequencing approaches in recovering coding DNA variation. Similarly, we performed this concordance analysis in the All of Us Research Program dataset to compare allele frequencies between whole genome sequencing and microarray genotyping to quantify any similar effect arising between these genotype discovery approaches in primarily non-coding variation.

We subsampled the All of Us primary release cohort down to the 95,596 samples who have both WGS and microarray genotyping data available. Call rate, HWE and MAF > 0.05 filters were applied to ensure only good quality common variants entered the analysis. Out of 102,631 variants (7,944 coding) -2,344 had Fisher’s exact test P < 0.05 (Supplemental File S1). Note that due to identical samples being analyzed, the expected p-value distribution is centered at 1; **Supplemental Figure S8**. We evaluated the overlap between the variants flagged in All of Us and gnomAD, finding that only 7 out of them were found in both samples, likely due to the focus of gnomAD on coding variation (given comparisons included WES) versus on non-coding variation in All of Us. Out of these 7 variants, rs4951250 was found to be significantly discordant in both datasets (P<1×10^−16^ genome vs exome; P=4×10^−5^ genome vs array).*Recovering filtered concordant sites*

In addition to generating this blacklist of bad sites that should be excluded despite being a PASS in gnomAD QC, we investigated whether additional trustworthy sites could be rescued from the ‘Non-PASS’ list based on our AF concordance criteria. Non-PASS variants are those that did not meet all required passing criteria in the gnomAD QC pipeline (Karczewski et al. 2019). We tested this by conditioning on PASS in one dataset, Non-PASS in the other, and re-ran the concordance pipeline requiring the following threshold in both datasets AC>1, DP > 10, and AF>0.01% to add a higher level of stringency for recovering sites. In total, there were 41,584 sites that met these criteria of which 30,683 were instances where the genomes are a Non-PASS and the exomes are a PASS. Approximately half of these sites had *p* values greater than 1×10^−5^, which we consider to be reliable. The exomes represent the vast majority of sequences in gnomAD, which may make their results more stable. For analyses that require less stringent QC, we provide these sites which can be optionally retained, given that they pass cohort QC in the individual dataset.

### Predicting technical bias for variants

We used features based on variant annotations generated during variant calling (e.g. variant quality, mapping quality, etc.) to build a random forest predictor that detects the presence of technical bias for a particular variant (**Supplemental Table S3**). Model training and validation was performed using several approaches. First, we used a leave-one-out cross validation procedure using the exome dataset from gnomAD. In 22 trials (one for each of the 22 autosomes) we set aside one chromosome and used the other 21 for model training. Then, the model was tested on the variants from the chromosome that was not used for training. Using the FET *p* value threshold for the discordance analysis we classified the variants into ‘bad’ (with *p* < threshold) and ‘good’ (*p* > = threshold) groups. By varying the FET *p* value threshold discriminating the groups we performed ROC analysis. Since representation of the classes varies depending on the selected threshold, we used the class weights for balancing the classes (in this and the following tests the weight for the ‘bad’ class was set as the fraction of ‘bad’ variants in the training data and the weight for the ‘good’ class was set to 1). The area under the ROC curve (ROC AUC) for such a model was estimated as 0.841 (**Figure 2A**).

**Figure 2.**
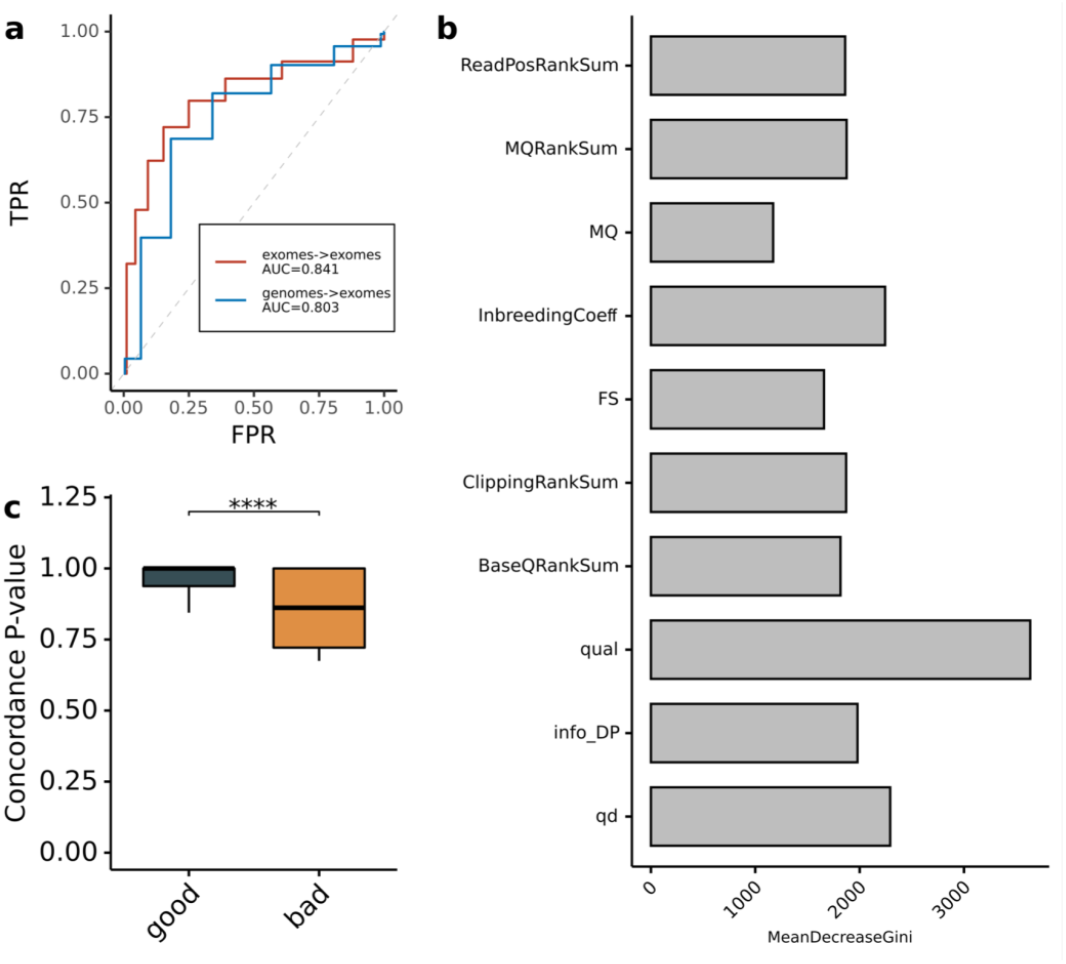
Reliable classification of discordant variants based on variant annotations. **A**) ROC analysis for two random forest predictor validation approaches using either leave-one-out analysis on the exomes, or the genomes as a training set to classify exome variants; **B**) Feature importance analysis for the random forest model; **C**) Comparison of concordance analysis FET *p* values for variants from 1000 Genomes classified using the random forest predictor trained on the gnomAD genomes dataset. MeanDecreaseGini is a measure of how each variable contributes to the homogeneity of the nodes and leaves in the resulting random forest. The higher the value of mean decrease accuracy or mean decrease Gini score, the higher the importance of the variable in the model.

Next, we used variant annotations from the gnomAD genomes dataset for training and variant annotations from the gnomAD exomes as a test sample. These are two separate datasets that are well-powered to detect biases in allele frequencies and ensure full independence between test and training samples. In this setting our model again reliably predicted discordant variants, with ROC AUC=0.803 (**Figure 2A**). Feature importance analysis of the model suggests that variant quality, inbreeding coefficient, and quality by depth are the key parameters discriminating variants with and without evidence for technical bias. Therefore, current protocols for alignment and variant calling are leaving a notable footprint which can be used to detect platform biases (**Figure 2B**).

Finally, we used data from the 1000 Genomes Project (1000 Genomes Project Consortium et al. 2012) as an independent public dataset for testing the predictor. Exome sequences from 1,393 samples (**Supplemental Table S4**) were used to create the variant call set following GATK best practices. Variant annotations were used to classify variants using the gnomAD genomes data as a training sample. Due to sample size, the 1000 Genomes data is significantly less powered to detect technical biases compared to gnomAD. Therefore, it is harder to confidently identify ground truth for discordant variants. Identical 1000 Genomes samples from exome and genome sequencing were used to detect variants with a signature of technical bias using the AF concordance FET described here (**Supplemental Table S5**). Due to the power limitations, instead of performing ROC analysis we compared the FET p values for the variants classified as having evidence of technical bias (‘bad’) and those without such an evidence (‘good’, **Fig 2C**).

This random forest model trained on the gnomAD genome v2.1 dataset was incorporated into a freely distributed R package called *DNAdiscover* -DNA DISCOrdant Variant IdentifiER (https://github.com/na89/DNAdiscover). The package uses variant annotations to predict whether a variant is likely to be ‘discordant’ or ‘concordant’ in user input data and performs well with both genome and exome sequencing data.

### Discordant variants are reported in published studies and are predicted to be functionally important

We first observed the phenomenon of variant discordance through the investigation of GWAS variants in the COVID host genetics initiative and the UK Biobank (COVID-19 Host Genetics Initiative 2021). When inspecting top associated variants that did not have strong LD friends, we noticed that many had discordant frequencies between GWAS arrays and gnomAD, and this often corresponded to variants that had discrepant frequencies between the gnomAD exomes and genomes. We thus suggest that the blacklist provided herein can be used for quality control of GWAS variants.

To investigate whether discordant variants may be spuriously attributed phenotypic relevance, we annotated all variants with their predicted functional consequence using the Ensembl Variant Effect Predictor (VEP) (McLaren et al. 2016). Blacklisted variants appear in all functional consequence categories, including 11536 that are annotated as missense (Supplemental Figure S9, Supplemental Table S5). Such variants are tempting to prioritize for functional follow-up given their apparent functional importance despite having GWAS signal likely driven by the observed genotype calling artifact. Technical artifacts are actually expected to be enriched in functionally important categories given that they are immune from the effects of natural selection, a phenomenon that has been previously observed for putatively loss-of-function somatic variation (Buckley et al. 2017).

To see if blacklist sites have been reported in peer-reviewed publications, we intersected genome-wide significant variants (*p* < 5 × 10^−8^) from the GWAS catalog (Welter et al. 2014) with our discordant sites list. Seventeen bad variants were found in the GWAS catalog, underscoring the importance of controlling for this artifact, as it may impact downstream interpretation of association findings (Supplemental Table S2). Variants in this list have been associated with multiple health-related phenotypes including schizophrenia, telomere length, and blood protein levels. Of these 17, half are multi-allelic and approximately a third are indels, echoing our earlier results of a higher discordance rate for these variant types than bi-allelic SNVs. Additionally, more than half of the 17 are present in the first megabase of the chromosome, suggesting that areas flanking the telomeres should be treated with caution.

## Discussion

The need for extremely large sample sizes to obtain sufficient statistical power in genetic studies requires the creation of datasets that may go beyond the financial capabilities of many individual research groups. This leads to the creation of meta-datasets that have contributions from many individual studies, thus creating heterogeneity in the genotype discovery approaches that were used for genotyping. Therefore, identification of DNA variants that are susceptible to technical bias when genotypes originate from multiple discovery strategies is vital in order to avoid false-positive associations and analyses of the artificially inflated allele frequencies. This is of particular concern in instances where cases may originate from one data generation effort and controls from another.

Here, we identify and describe a technical artifact arising in various genotype discovery approaches that may affect cohort data variant quality despite the following of gold standard QC procedures. We present our metric for the identification of discordant sites, provide a blacklist of the discordant variants identified in gnomAD which should be treated with caution, and release an openly-available software package containing our random forest predictor that reliably classifies untrustworthy variants in user cohort data. Excluding variants with signals of discordance across sequencing platforms will result in higher quality results and reduces the risk of spurious associations in gene discovery. This is particularly important as we observe that technical artifacts are enriched in functionally important annotations.

Additionally, we show that discordance in allele frequencies is also present in the All of Us Research Program dataset, when comparing whole genome sequencing to microarray genotyping for overlapping samples. This finding indicates that variants in both coding and non-coding DNA could have discordant genotype calls. Importantly, in our predictor we use the variant annotations which often are utilized in variant quality score recalibration and filtration pipelines. Our results indicate that stricter filtration thresholds might be helpful for elimination of some discordant variants, however, more cautious consideration of discordance is warranted in heterogeneous datasets.

We note that while we provide a blacklist of variants failing our discordance test in the gnomaD v2 dataset for ready exclusion, the specific sites that are discordant in a given cohort depends on the genotype discovery approach utilized and dataset composition. Therefore, for optimal precision, we recommend identification of discordant sites within user cohorts with the provided classifier rather than a blanket restriction of variants identified in gnomAD. We freely provide an R-package with a predictor trained on gnomAD whole genome sequencing data, *DNAdiscover*, for such use in other cohort data to identify cohort-specific sites with features indicative of unreliable genotype calls.

Based on the examinations presented in this manuscript, we recommend researchers using aggregated cohort data to implement the following conservative QC procedures to ensure the elimination of discordant sites:

- Drop any variant that fails in both the gnomAD exomes and genomes
- Consider dropping any variant that fails in the gnomAD exomes, as these represent the bulk of gnomAD data
- Drop the blacklist variants presented here that are PASS in both the gnomAD exomes and genomes but that are discordant in frequency across genotype discovery approach
- Drop variants that are flagged by our random forest predictor, *DNAdiscover*, in an independent dataset, as each genotype discovery approach has a distinct genotyping error mode
- Remove the low complexity regions
- Optional: skeptically retain sites that are on the ‘recovered’ list here

## Methods

### Characterizing discordance in genotype calls across gnomAD exomes and genomes

All analyses were conducted using the Hail software program on the Google Cloud platform (GCP 2021). Plots were created using Bokeh and ggplot2 (Jolly 2018; Wickham 2011). Concordance metrics for genotype calls were generated from the overlapping individuals with the command hail.methods.concordance(). Using the full release of gnomAD version 2.1 (Karczewski et al. 2020), we filtered to include only sites that were both present and had a quality determination of PASS in the genomes and exomes. We split multi-allelic variants, retained only sites that are present and PASS in both exomes and genomes, filtering to only bi-allelic sites with AC > 1 and AF > 0.01% in either dataset for the NFE. Starting with all sites with at least 1 alternate allele, and subsequently for sites with AC>5 and 10, we calculated the allele frequency (AF) in the genomes and exomes separately and ran a Fisher’s Exact Test (FET) on the difference in the number of alternate AC to total alleles (allele number, AN) between these two datasets. Specific filters for various steps are described in their relevant *Results* section.

AllofUs data was subsampled to 95,596 with both whole genome sequencing (WGS) and microarray genotyping (WGA) available. MAF>0.05, HWE > 0.0001 and MAC>10 filters were applied to keep only common variants. Multiallelic variants were split. Variant call rate > 0.8 was required in both WGS and WGA datasets. Variants with call rate difference between datasets greater that 0.05 were also eliminated from analysis. The R libraries *dplyr, reshape2, pROC, ROCR* and *RandomForest* were used to process variant annotations, evaluate predictor quality and build a DNAdiscover using R-4.0.3 (Wickham et al. 2015; Wickham 2012; Robin et al. 2011; Sing et al. 2005; Liaw et al. 2002).

## Acknowledgements

We thank the members of the gnomAD and Hail teams for their assistance with this project. This project was supported by the National Institute of Mental Health (K01 MH121659 and T32 MH017119 to E.G.A.) and the National Human Genome Research Institute (U24HG011450). E.G.A was additionally supported by the Caroline Wiess Law Fund for Research in Molecular Medicine and the ARCO Foundation Young Teacher-Investigator Fund at Baylor College of Medicine. A.A.L. was supported by the Ministry of Science and Higher Education of the Russian Federation (Agreement No. 075-15-2022-301, Institutional grant to Almazov National Medical Research Center). M.A. was supported by the Aging Biology Foundation and Nationwide Foundation Pediatric Innovation Fund. The All of Us Research Program is supported by the National Institutes of Health, Office of the Director: Regional Medical Centers: 1 OT2 OD026549; 1 OT2 OD026554; 1 OT2 OD026557; 1 OT2 OD026556; 1 OT2 OD026550; 1 OT2 OD 026552; 1 OT2 OD026553; 1 OT2 OD026548; 1 OT2 OD026551; 1 OT2 OD026555; IAA #: AOD 16037; Federally Qualified Health Centers: HHSN 263201600085U; Data and Research Center: 5 U2C OD023196; Biobank: 1 U24 OD023121; The Participant Center: U24 OD023176; Participant Technology Systems Center: 1 U24 OD023163; Communications and Engagement: 3 OT2 OD023205; 3 OT2 OD023206; and Community Partners: 1 OT2 OD025277; 3 OT2 OD025315; 1 OT2 OD025337; 1 OT2 OD025276. In addition, the All of Us Research Program would not be possible without the partnership of its participants.

## Author Contributions

E.G.A. and M.A. designed and implemented pipelines, ran analyses, and drafted the primary manuscript. K.J.K., A.A.L. aided in code implementation and interpretation. H.R., D.M., B.M.N. and M.J.D. supervised and advised on the project. All authors reviewed and approved the final draft.

## Data Access

gnomAD summary data is freely available at https://gnomad.broadinstitute.org/. Additional information and a discussion of the best practices for using gnomAD can be found at https://macarthurlab.org/blog/. *DNAdiscover* package for prediction of presence of the technical bias in variants coming from NGS and usage manual are available at https://github.com/na89/DNAdiscover. A blacklist of bad variants failing the discordance test in gnomAD is provided with this manuscript in the Supplementary Materials.

