## Supplemental Information for "Discordant calls across genotype discovery approaches elucidate variants with systematic errors"

### Supplemental Figures

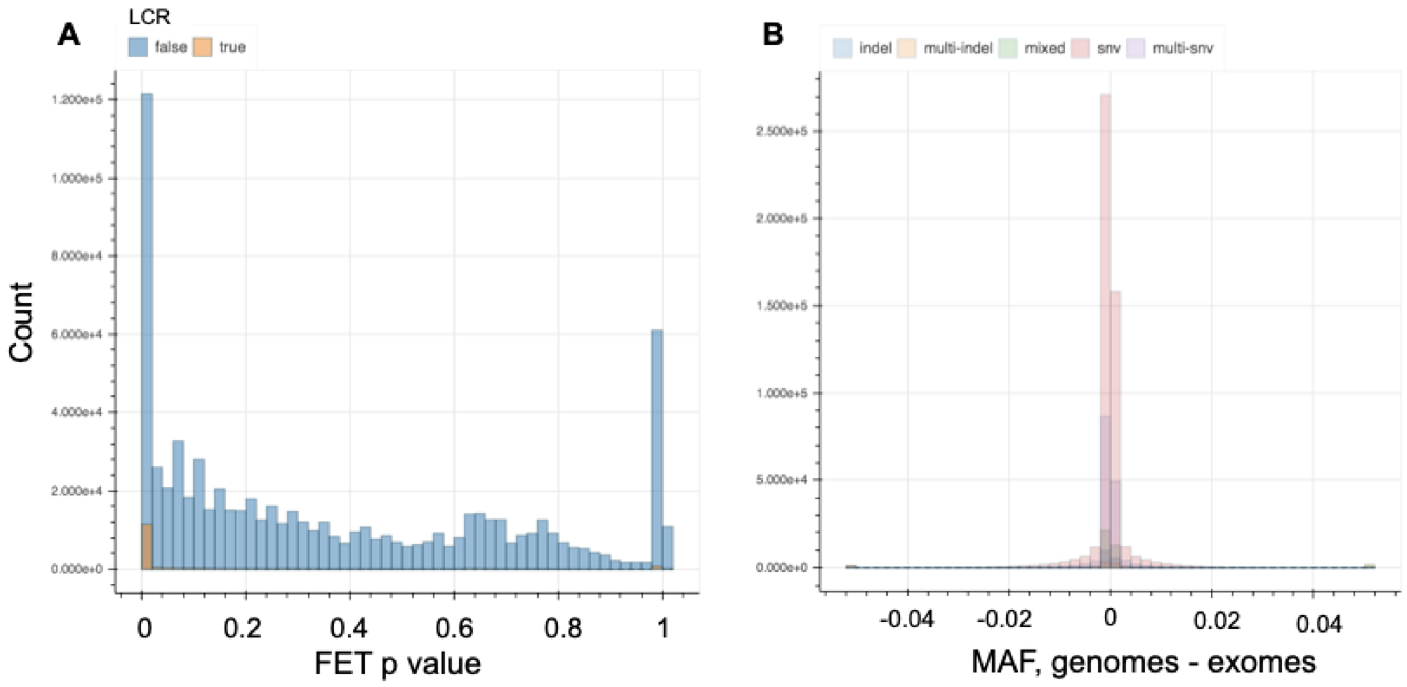


**Supplemental Figure S1. The signal of discordance is consistent across AC thresholds. (A)** FET p value for variants with at AC>1. Variants falling in the low complexity region (LCR) are indicated with orange and are enriched in the worst performing bin. **(B)** Distribution of allele frequencies in the NFE exomes vs genomes at AC>5. Variants are colored by variant type. Both panels show results for the NFE.

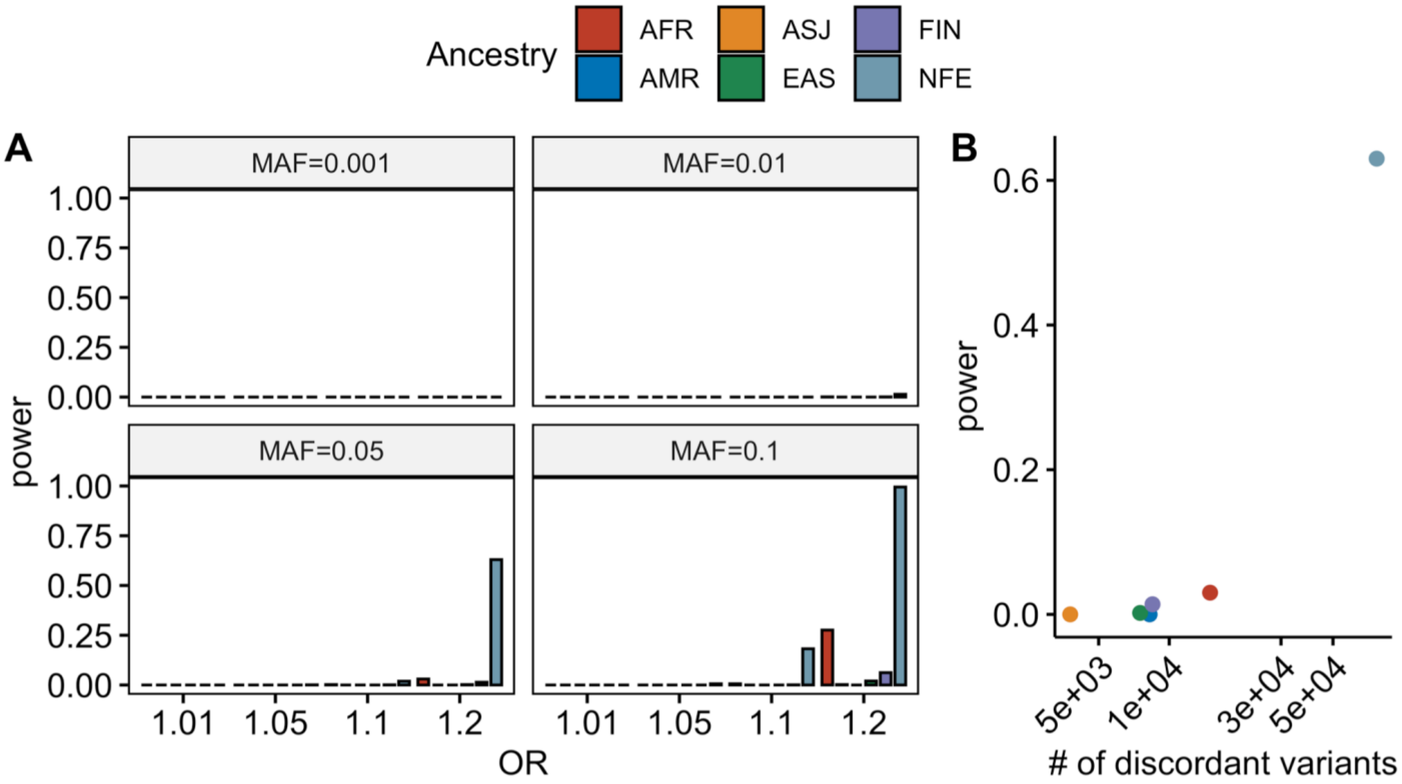


**Supplemental Figure S2. Power to detect variants with discordant allele frequencies between genotype discovery approaches. (A)** Fisher's exact test power evaluated for the number of samples in gnomAD WES and WGS datasets for each ancestry given a particular MAF cutoff; **(B)** Fisher's exact test power for MAF=0.05 and OR=1.2 and the observed number of discordant variants ($p<1x10^{-5}$) for each ancestry. Note that due to sample size the NFE are the most highly powered for identifying discordant variants.

**A**

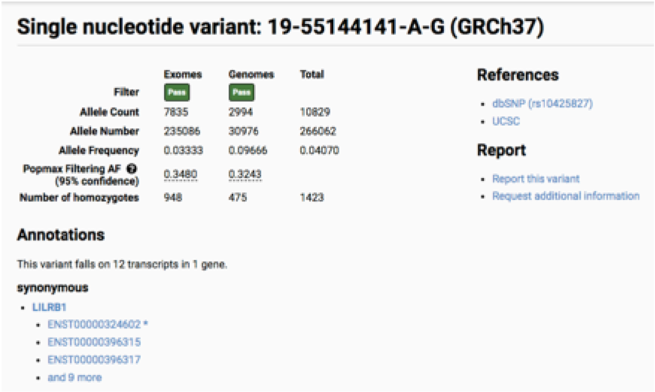


**B**

| | 0 | 1 | 2 | 3 | 4 |
|---|---|---|---|---|---|
| **0** | Missing variant both | Missing variant G | Missing variant G | Missing variant G | Missing variant G |
| **1** | Missing variant E | No call both | No call G | No call G | No call G |
| **2** | Missing variant E | No call E | ref/ref both | Hom ref G / Het E | Hom ref G / Hom Var E |
| **3** | Missing variant E | No call E | Het G / Hom ref E | het both | Het G / Hom var E |
| **4** | Missing variant E | No call E | Hom var G / hom ref E | Hom var G / het E | hom var both |

**Supplemental Figure S3. Examples of discordance. (A)** Example of a discordant variant as seen in the gnomAD browser. Note that this variant is PASS in both the Exomes and Genomes, but that there is a sizable MAF difference depending on genotype discovery approach. **(B)** Concordance table. The miscall categories considered as discordant here are shown in white. Gray indicates variants that were excluded from the concordance test due to missing information in one of the two datasets. Red indicates no alternative alleles were observed in either dataset. Green indicates concordant calls when alternative alleles were observed.

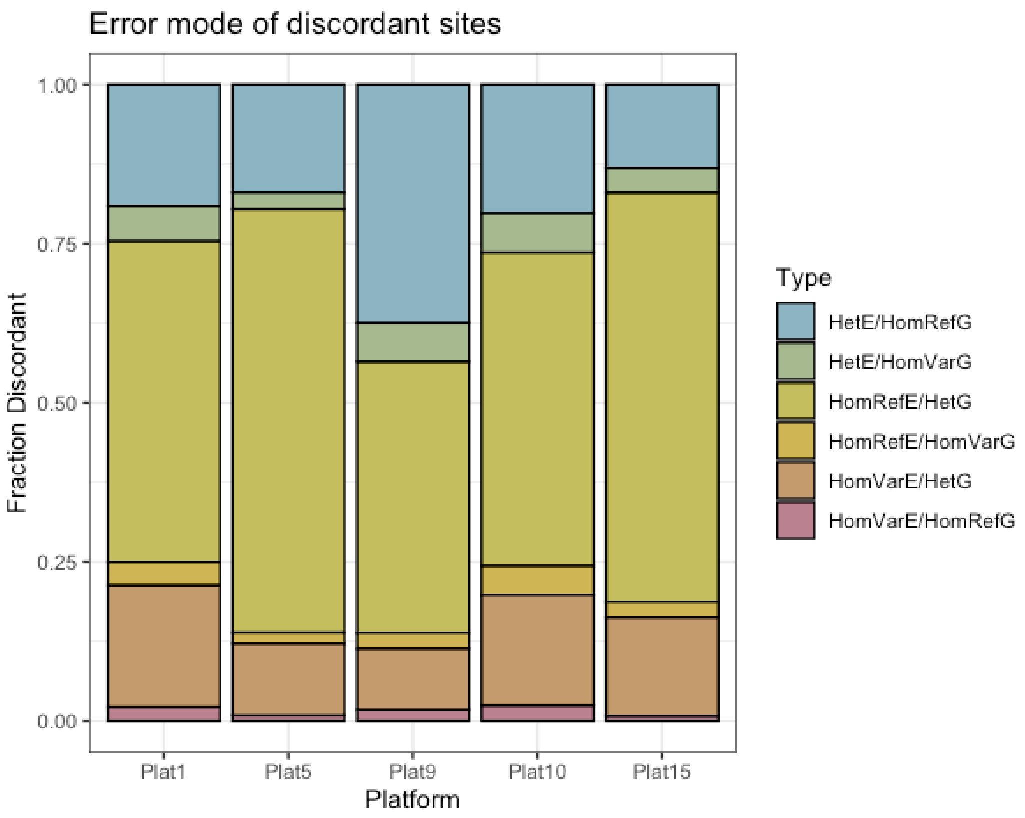


**Supplemental Figure S4. Wrong call error mode by technology platform**. The proportion of wrong calls in each error mode category are shown for each gnomAD sequencing platform for which overlapping data was available.

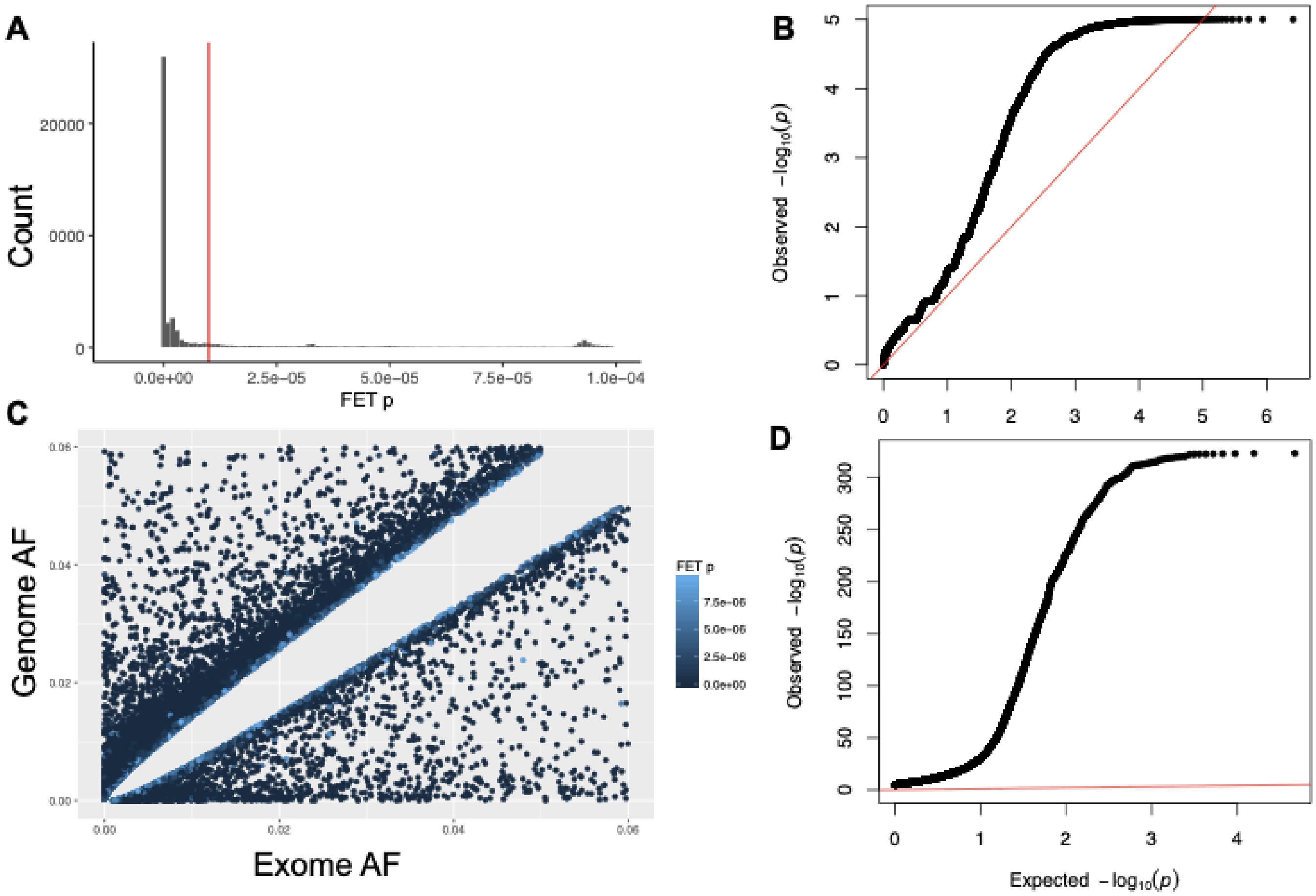


**Supplemental Figure S5. Selection of 1e-5 as the threshold for 'good' vs 'bad' variants. (A)** Concordance FET *p* value for AC>10 highlighting 1e-5, indicated with the red line, which was chosen as the threshold for NFE variants considered to be discordant. **(B)** QQ plot for the concordance test for all NFE variants. **(C)** Exomes vs genomes AF for bad NFE variants failing the 1e-5 threshold. **(D)** QQ plot for just the bad variants.

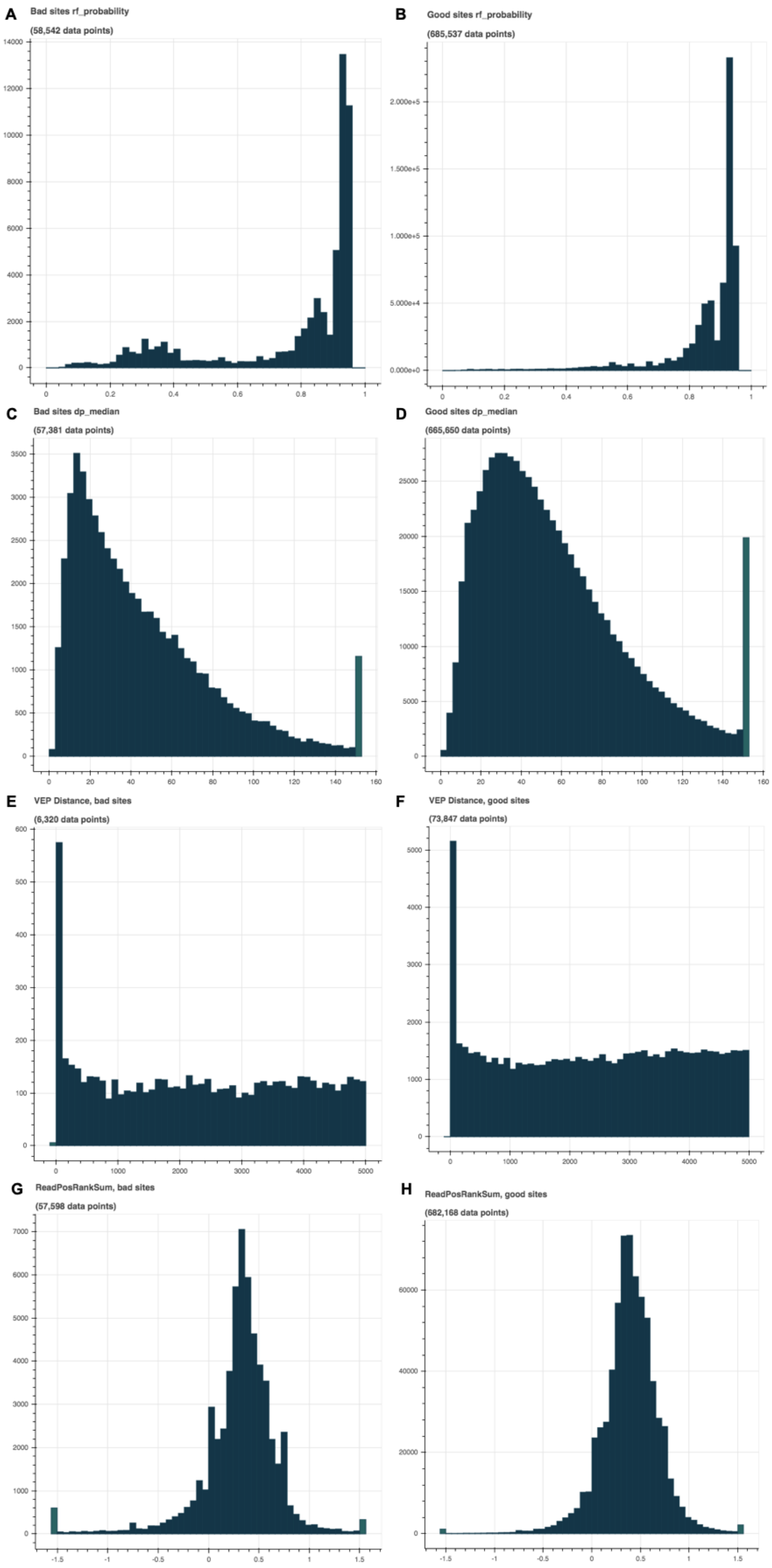
A
Bad sites rf_probability
(58,542 data points)
B
Good sites rf_probability
(685,537 data points)
C
Bad sites dp_median
(57,381 data points)
D
Good sites dp_median
(665,650 data points)
E
VEP Distance, bad sites
(6,320 data points)
F
VEP Distance, good sites
(73,847 data points)
G
ReadPosRankSum, bad sites
(57,598 data points)
H
ReadPosRankSum, good sites
(682,168 data points)

**Supplemental Figure S6**. Distribution of gnomAD exomes metadata features for good versus bad variants. Bad variant distributions are presented on the left, good on the right. **(A,B)** RF_Probability, the confidence of the random forest genotyper implemented with gnomAD. **(C,D)** DP_Median, the median depth of exomes. **(E,F)** VEP Distance, the distance to the closest canonical gene. **(G,H)** ReadPosRankSum, how far along sequencing reads the variant is falling.

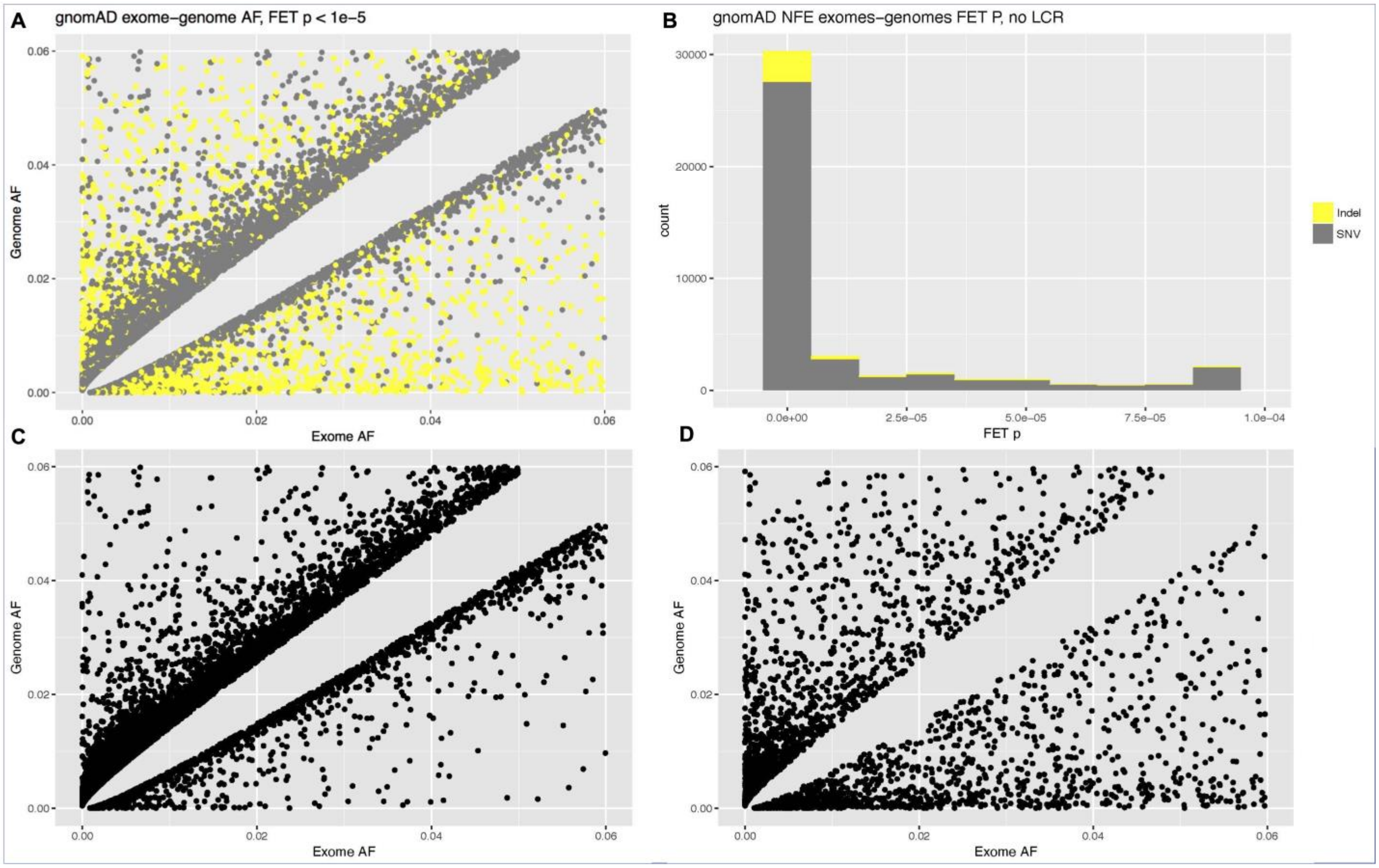

**Supplemental Figure S7. Different patterns of discordance in SNVs as compared to indels. (A) (B)** FET *p* value for variants after excluding the LCR region. **(C)** Exomes vs genomes AF for bad NFE SNV variants failing the 1e-5 threshold. **(D)** Exomes vs genomes AF for indel bad NFE indels failing the 1e-5 threshold.

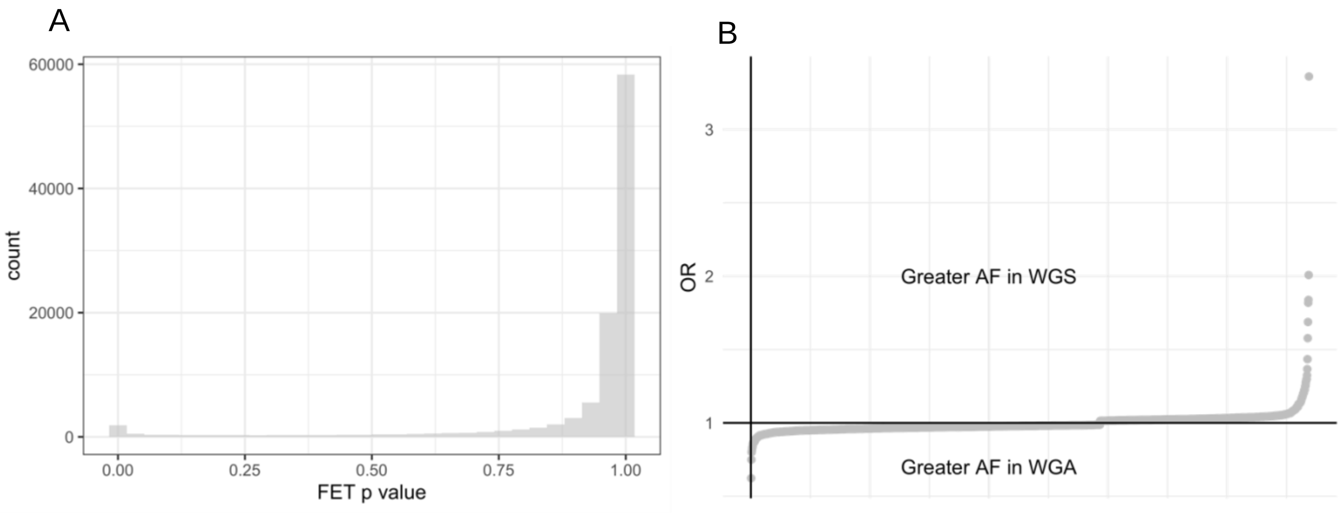


**Supplemental Figure S8. Analysis of variant allele frequency concordance between whole genome sequencing and microarray data in the All of Us Research Program.** (A) Distribution of FET p-values for the allele frequency comparison between the whole genome sequencing and microarray genotyping for the same individuals from All of Us. (B) Odds ratios for the variants with FET $p < 0.05$.

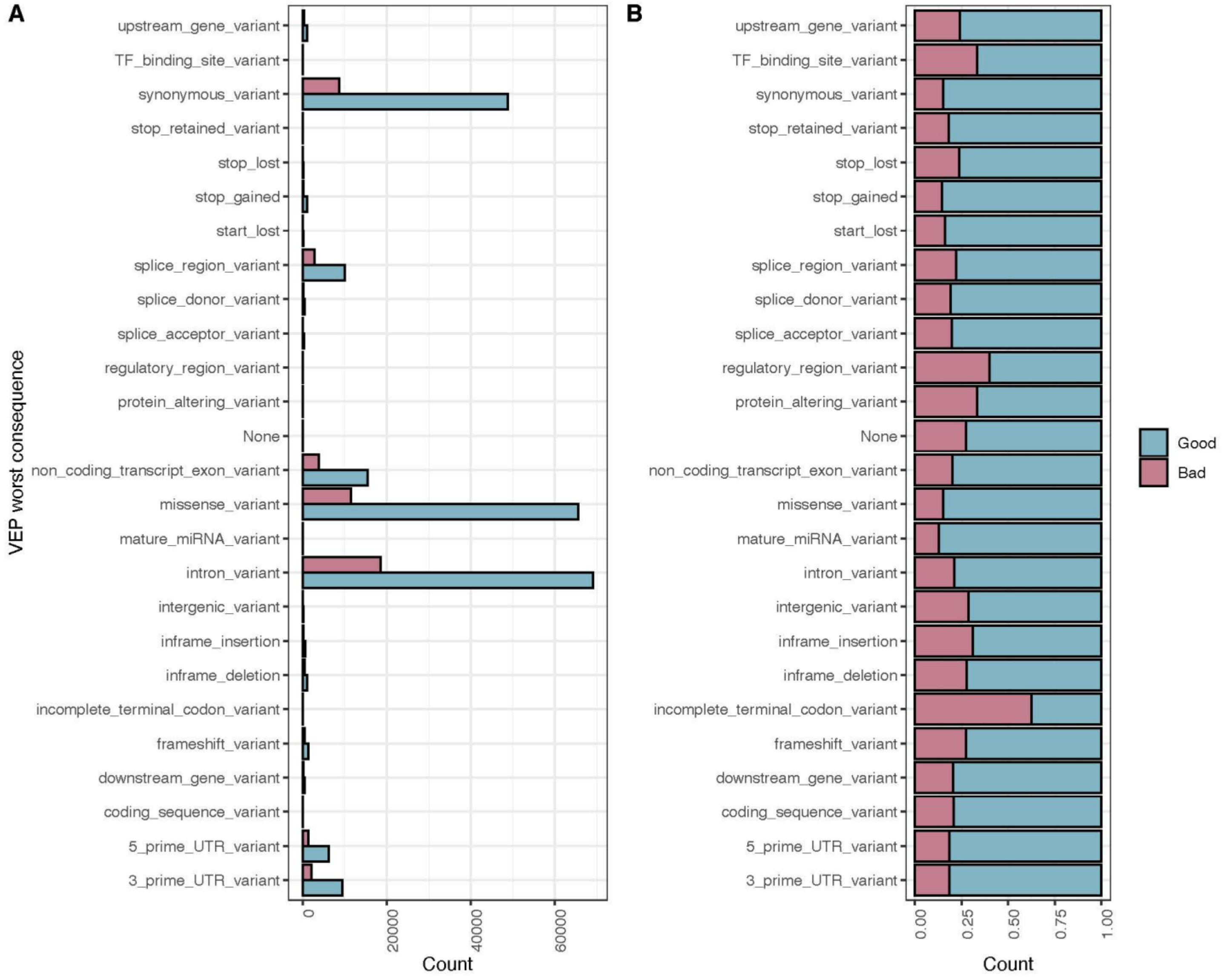


**Supplemental Figure S9**. **VEP predicted worst consequence for ‘good’ versus ‘bad’ variants with MAF>0.01 after an AC > 10 filter.** Note the presence of many bad variants that are predicted to have severe functional consequences. (A) Absolute count; (B) proportion of total in each category.

### Supplemental Tables

| **Population** | **Fraction bad sites shared with NFE** |
|---|---|
| AFR | 0.745 |
| AMR | 0.865 |
| ASJ | 0.992 |
| EAS | 0.803 |
| FIN | 0.938 |
| **Average** | **0.869** |

**Supplemental Table S1.** Large overlap in discordant sites between NFE and other continental ancestry groups.

| Chromosome | Position |
|---|---|
| 4 | 3494956 |
| 5 | 33963745 |
| 6 | 31852866 |
| 6 | 151687847 |
| 9 | 712766 |
| 9 | 4576774 |
| 9 | 5126343 |
| 9 | 5185581 |
| 11 | 308180 |
| 11 | 308290 |
| 11 | 308314 |
| 11 | 309127 |
| 11 | 828916 |
| 18 | 618124 |
| 18 | 662103 |
| 19 | 844020 |
| 19 | 913048 |

**Supplemental Table S2.** Discordant variants seen in the GWAS catalog.

| |
|---|
| Low complexity region membership |
| segdup |
| nonpar |
| variant_type |
| allele_type |
| was_mixed |
| has_star |
| qd |
| info_SOR |
| rf_probability |
| was_split |
| score |
| qual |
| BaseQRankSum |
| ClippingRankSum |
| FS |
| InbreedingCoeff |
| MQ |
| MQRankSum |
| ReadPosRankSum |

**Supplemental Table S3.** Features of the gnomADv2 variant annotations used in the random forest prediction model.

| **Ancestry** | **AFR** | **AMR** | **EAS** | **EUR** | **SAS** |
|---|---|---|---|---|---|
| **Number of Samples** | 296 | 222 | 422 | 351 | 102 |

**Supplemental Table S4.** 1000 Genomes samples per ancestry included into testing dataset.

| **VEP_worst_consequence** | **All** | **Bad** | **Good** |
|---|---|---|---|
| 3_prime_UTR_variant | 11519 | 2130 | 9380 |
| 5_prime_UTR_variant | 7602 | 1399 | 6175 |
| coding_sequence_variant | 35 | 7 | 27 |
| downstream_gene_variant | 596 | 121 | 475 |
| frameshift_variant | 1833 | 494 | 1312 |
| incomplete_terminal_codon_variant | 8 | 5 | 3 |
| inframe_deletion | 1485 | 410 | 1075 |
| inframe_insertion | 791 | 244 | 547 |
| intergenic_variant | 263 | 75 | 188 |
| intron_variant | 87799 | 18589 | 69116 |
| mature_miRNA_variant | 95 | 12 | 83 |
| missense_variant | 77243 | 11536 | 65588 |
| non_coding_transcript_exon_variant | 19353 | 3858 | 15463 |
| regulatory_region_variant | 50 | 20 | 30 |
| splice_acceptor_variant | 467 | 91 | 370 |
| splice_donor_variant | 554 | 106 | 446 |
| splice_region_variant | 12916 | 2842 | 10053 |
| start_lost | 277 | 44 | 232 |
| stop_gained | 1257 | 180 | 1077 |
| astop_lost | 195 | 46 | 148 |
| stop_retained_variant | 72 | 13 | 59 |
| synonymous_variant | 57514 | 8703 | 48755 |
| TF_binding_site_variant | 3 | 1 | 2 |
| upstream_gene_variant | 1341 | 323 | 1013 |
| None | 10 | 3 | 8 |
| protein_altering_variant | 9 | 3 | 6 |

**Supplemental Table S5**. **Counts for good and bad variants within VEP categories (AC >10)**.

Note the substantial numbers of bad variants with severe predicted consequences.
